# Mechanism and consequence of daily modulation of cortical parvalbumin-positive inhibitory neurons

**DOI:** 10.1101/2022.03.30.486351

**Authors:** Fang-Jiao Zong, Xue-Ting Zhang, Yan Zhang, Xia Min, Yang Liu, Kai-Wen He

## Abstract

Parvalbumin-positive (PV) neurons, the main class of inhibitory neurons in the neocortex, play critical roles in maintaining normal brain function and are implicated in a variety of brain disorders. Here we found that their function is modulated in a time- and sleep-dependent manner naturally during the day. We first show that PV-evoked inhibition is stronger by the end of the light (ZT12) compared to the end of dark (ZT0) cycle. In addition, both PV’s excitatory and inhibitory synaptic transmission slowly oscillate but in the opposite directions during the light/dark cycle. Whereas excitatory synapses are predominantly regulated by experience, inhibitory synapses are regulated by sleep. Mechanistically, we found that the daily regulation of PV’s inhibitory synapses is mediated by acetylcholine activating M1 receptors. Consistent with our ex vivo findings, we show in vivo that PV’s spontaneous activity display clear oscillation, which is opposite to that of the pyramidal neurons. Finally, we demonstrate that the daily changes in PV neural activity negatively correlate with the dLGN-evoked responses in V1, underscoring the physiological significance of PV’s daily regulation.

## Introduction

Parvalbumin (PV)-expressing GABA (ɣ-aminobutyric acid)-ergic neurons are the most abundant class of inhibitory neurons in the neocortex(Rudy, Fishell et al., 2011). By forming dense inhibitory connections around soma or the axon initial segment of their neighboring neurons, mainly glutamatergic principal neurons, these fast-spiking inhibitory neurons gate the excitability of their targets and play critical roles in maintaining the stability as well as enhancing the computational power and precision of the neural network(Ferguson & Cardin, 2020, Hu, Gan et al., 2014, Sadeh & Clopath, 2021). For example, in the primary visual cortex (V1), PV neurons are known to control cortical plasticity(Sur, Nagakura et al., 2013, van Versendaal & Levelt, 2016) and visual response properties such as response gain(Atallah, Bruns et al., 2012, Wilson, Runyan et al., 2012, Zhu, Qiao et al., 2015). Furthermore, dysregulation of PV neuron is tightly associated with a variety of brain disorders, such as epilepsy(Wang, Xu et al., 2017), autism(Vogt, Cho et al., 2018), schizophrenia(Steullet, Cabungcal et al., 2017) and Alzheimer’s disease(Hijazi, Heistek et al., 2019). Thus, defining the functional regulation of PV neurons will provide important insight into their role in normal and diseased brains.

Despite their functional importance, how PV neurons are modulated remains poorly understood. Most studies have focused on the impacts of disease-associated genetic mutations on early development of PV neurons(Hu, Vogt et al., 2017, Marin, 2012), which provides insight into the cause of defects in these diseases. But for other psychiatric and neurodegenerative disorders, very little is known about how PV neuron dysregulation might occur. Recent studies showed that early-life sleep may regulate the neurodevelopment of cortical PV neurons(Jones, Opel et al., 2019). In mature brain, different sleep/wake stages and sleep history are reported to acutely affect the spontaneous firing of PV(Niethard, Hasegawa et al., 2016) or general inhibitory neurons(Miyawaki & Diba, 2016, Vyazovskiy, Olcese et al., 2009), although contradictory results exist(Hengen, Lambo et al., 2013, Hengen, Torrado Pacheco et al., 2016). These findings suggest sleep may be crucial for PV regulation. We previously reported that the excitation/inhibition (E/I) ratio of cortical pyramidal neurons naturally changes during the 24-hour day, of which the synaptic inhibition oscillates dramatically and is influenced by sleep(Bridi, Zong et al., 2020). Taken together, we hypothesized that cortical PV neurons might be dynamically modulated daily. Whether and how this may happen and what physiological impacts it may have are questions of importance that may shed light into the functional modulation of PV neurons.

We studied the above questions in V1 by combining ex vivo and in vivo assessments. We first observed that the inhibitory output from PV neurons indeed dramatically differs between the light and dark cycle. Consistently, we found that PV’s excitatory and inhibitory synaptic transmission slowly oscillate in the opposite direction during the 24-hour day. In addition to the time-dependency, we found that experience dominates the regulation of PV’s synaptic excitation, while sleep plays essential role in regulating PV’s synaptic inhibition. We further identified ACh via targeting presynaptic M1R is required for the daily regulation of synaptic inhibition. In line with all the *ex vivo* changes, we observed a daily oscillation in PV spontaneous activity *in vivo*. The activity of putative pyramidal neurons in V1 negatively correlates with PV’s. Finally, we demonstrated that direct dorsal lateral geniculate nucleus (dLGN) stimulation evoked neuronal activities in V1, which mimics vision-evoked responses, are stronger at ZT0 compared to ZT12. This pattern is consistent with PV’s role in gain control, further strengthening the physiological importance of this novel mode of daily modulation of PV function.

## Results

### Altered PV neuron-evoked inhibitory output between the light and dark cycle

We previously found that the inhibitory synaptic transmission of V1 L2/3 pyramidal neurons slowly increases and remains high during the light cycle and gradually reduces and remains low during the dark cycle(Bridi et al., 2020). Therefore, we hypothesized that altered inhibitory output from PV neurons contribute to this phenomenon. By using PV:Ai32 transgenic mice expressing channelrhodopsin-2 (ChR2) specifically in PV neurons, we directly measured light-evoked inhibitory postsynaptic currents (IPSCs) in V1 L2/3 pyramidal neurons at the end of the light cycle (Zeitgeber time 12, ZT12) and the end of dark cycle (ZT0) (Fig 1A & Fig S1A, see Methods). In V1-containing acute brain slice, we sub-divided a cubic area of 0.45 mm2 covering L1 to L5 of V1 into 15 by 15 grids (Fig 1A_3_). The soma position of the recorded neurons was relatively fixed. Each grid was sequentially stimulated with brief blue light pulse in a pseudorandom order and pharmacologically isolated, light-evoked inhibitory postsynaptic currents (_LE_IPSCs) were recorded from pyramidal neurons voltage clamped at -50 mV. We only analyzed _LE_IPSCs with short latency, high fidelity and maximal amplitude larger than 60 pA (see Methods). We used the maximal amplitude of the _LE_IPSCs as pixel value for corresponding grid and plotted a 2-dimensional map for each pyramidal neuron (Fig 1A_5_, see Methods). We first compared the IPSC strength by aligning all maps from each group according to the soma position of the recorded neuron. The averaged soma-aligned map of ZT12 shows higher signal intensity than ZT0 (Fig 1B). _LE_IPSCs at ZT12 were significantly larger (both max amplitude and summation of all currents) compared to those at ZT0 (Fig 1C-D). The difference between ZT0 and ZT12 was unlikely caused by variant levels of ChR2 since we randomly recorded PV:Ai32 mice from the same litter at ZT0 or ZT12, and the differences were highly conserved among all litters (Fig S1B). These results support our hypothesis that the functional output from cortical PV neurons varies at different times of the day – high at ZT12 while low at ZT0.

**Figure 1.**
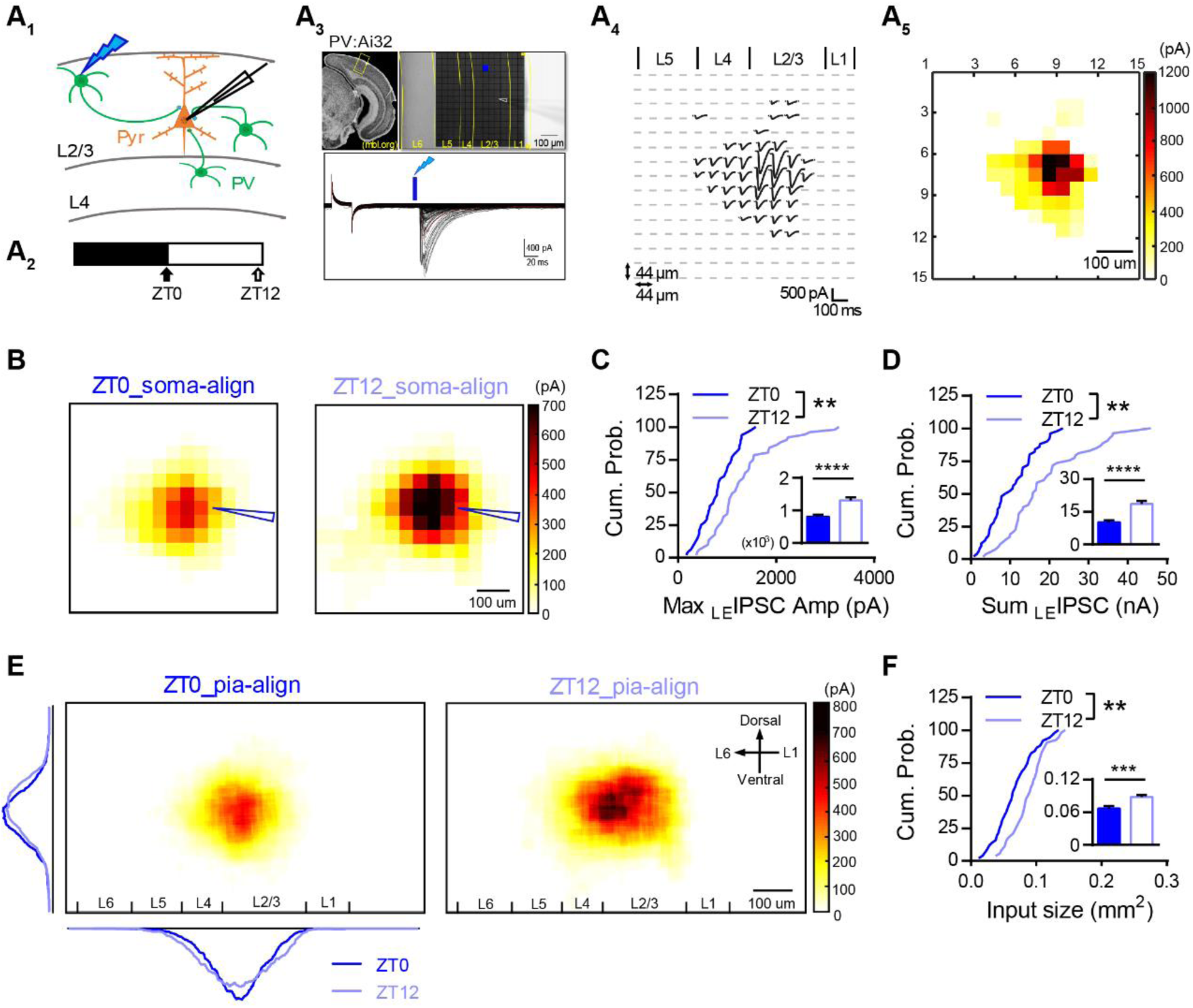
Distinct inhibitory output from PV neurons between the light and dark cycle. A. Experimental demonstration. A_1_. Illustration of the mapping principle between ChR2-expressed PV neurons (green) and regular pyramidal neuron (Pyr, orange). A_2_. Acute slices were obtained and recorded at ZT0 or ZT12. A_3_. Coronal brain atlas showing the primary visual cortex (Top left panel, modified from MBL mouse brain atlas). Yellow box highlights the region of experiment. Top right panel, the reference image taken with a 10x objective from a slice prepared from a PV:Ai32 mouse. The 0.45 mm^2^ shaded cubic area was divided into 15 × 15 grids for independent light stimulation. The white arrowhead indicates the position of the recording electrode. Bottom panel, representative raw traces of light-evoked IPSCs (_LE_IPSC) recorded from one pyramidal neuron. A_3_. The spatial distribution of the _LE_IPSCs from the example in A_2_. A_4_. The heatmap of the _LE_IPSCs in A_3_. B. The average heatmaps at ZT0 and ZT12 after soma alignment. Soma position was indicated by the blue arrow head. C. Comparison of the cumulative distribution (*K-S* test, P = 0.002) and the mean (Insert. ZT0: 813.3 ± 53.13 pA; ZT12: 1306 ± 93.73 pA; P < 0.0001) of the max _LE_IPSC amplitude between ZT0 and ZT12. D. Comparison of the cumulative distribution (*K-S* test, P = 0.0016) and the mean (Insert. ZT0: 10.44 ± 1.44 nA; ZT12: 18.87 ± 1.44 nA; P < 0.0001) of the sum _LE_IPSC amplitude between ZT0 and ZT12. E. The average heatmaps of _LE_IPSCs after pia aligned to the reference image in A_2_. Dark and light blue overlay traces on the side of the map depicted the marginal distribution of the sum _LE_IPSCs for ZT0 (Dark blue) and ZT12 (Light blue). F. Comparison of the cumulative distribution (*K-S* test, P = 0.0021) and the mean (Insert, ZT0: 0.067 ± 0.004 mm^2^; ZT12: 0.089 ± 0.0036 mm^2^; P = 0.0003) inhibitory input sizes between ZT0 than ZT12. Sample sizes used in this figure. ZT0: 48 cells from 9 mice; ZT12: 51 cells from 9 mice.

To further investigate whether the spatial distribution of PV-evoked inhibition was also altered between ZT0 and ZT12, we projected each map onto the reference map (Fig 1A_3_) after pia alignment (pia-align, see Method). The averaged pia-aligned maps of both groups show that the _LE_IPSCs were mainly evoked within L2/3 with similar lateral distribution between groups (Fig 1E). In the vertical direction, a significant proportion of IPSCs was also evoked from L4, but relatively weak from L5, and the distribution seems to be flattened at ZT12 (Fig 1E). While pyramidal neurons recorded in both groups had similar spatial distribution (Fig S1C-D), their input size was significantly larger at ZT12 than at ZT0 (Fig 1F), suggesting PV neurons from broader area might be recruited at ZT12. Thus, these results in combine suggest that PV output indeed alters markedly at different times of the day.

### Opposite oscillation of excitatory and inhibitory synaptic transmission of PV neurons during the light/dark cycle

The balance between synaptic excitation (E) and inhibition (I) could dictate somatic firing(Gidon & Segev, 2012, Liu, 2004). We previously found that E and I of cortical pyramidal neurons are modulated daily(Bridi et al., 2020). We therefore wondered whether E and I of PV neurons are also modulated during the light/dark cycle. We recorded both miniature excitatory postsynaptic currents (mEPSCs) and inhibitory postsynaptic currents (mIPSCs) of V1 L2/3 PV neurons from acute brain slices harvested at either ZT0 or ZT12. mEPSC frequency was higher at ZT12 than at ZT0 (Fig 2A), suggesting that PV neurons receive more excitatory inputs by the end of the light cycle. Interestingly, mIPSC frequency was significantly reduced at ZT12 compared to ZT0 (Fig 2B), which indicates PV neurons are less suppressed at ZT12. There was no change in the amplitude of mEPSCs and mIPSCs (Fig 2C-D), suggesting a presynaptic rather than postsynaptic locus for the altered synaptic transmission.

**Figure 2.**
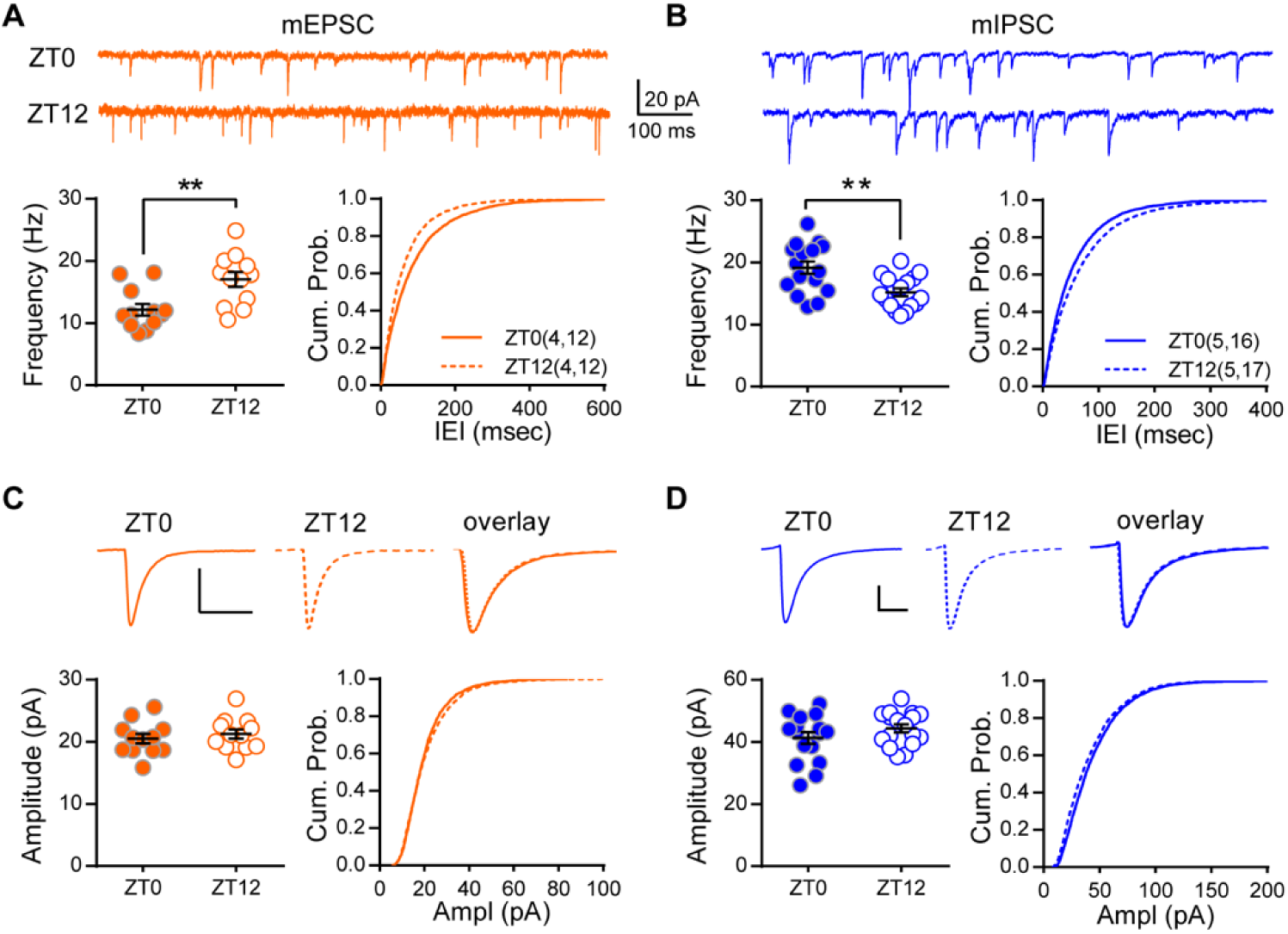
Opposite modulation of PV’s synaptic excitation and inhibition during the light/dark cycle. A. mEPSC frequency was higher at ZT12 than ZT0. Top: example traces of mEPSC recordings. Bottom left: mEPSC frequency (ZT0: 12.17 ± 0.94Hz; ZT12: 17.03 ± 1.20Hz). Bottom right: cumulative probability histogram of the inter-event intervals (*K-S* test, P < 0.0001). B. mIPSC frequency was lower at ZT12 than ZT0. Top: example traces. Bottom left: mIPSC frequency (ZT0: 19.15 ± 0.99Hz; ZT12: 15.19 ± 0.63Hz). Bottom right: cumulative probability histogram of the inter-event intervals of mIPSCs (*K-S* test, P < 0.0001). C. mEPSC amplitude and kinetics did not differ between ZT0 and ZT12. Top: average traces of the well-isolated events. Bottom left: mEPSC amplitude (ZT0: 20.51 ± 0.78pA; ZT12: 21.26 ± 0.75pA; P = 0.34). Bottom right: The distribution of mEPSC amplitude of all events (*K-S* test, P < 0.01). D. mIPSC amplitude and kinetics did not differ between ZT0 and ZT12. Top: average traces of the well-isolated mIPSCs. Bottom left: mIPSC amplitude (ZT0: 41.31 ± 1.91pA; ZT12: 44.39 ± 1.31pA; P = 0.35). Bottom right: The cumulative distribution of mIPSC amplitudes (*K-S* test, P < 0.0001). Sample size is indicated as (mice, cells).

To further evaluate the temporal profile of synaptic transmission, we recorded spontaneous EPSCs and IPSCs that are more physiologically relevant(Bridi et al., 2020, Jurgensen & Castillo, 2015) at multiple time points during the light/dark cycle (Fig 3A). Similar to miniature currents, both sEPSCs and sIPSCs changed but in the opposite direction. Specifically, sEPSCs and sIPSCs were increased and decreased respectively 4 hours after entering the light cycle, then they were maintained relatively stable between ZT4 and ZT12. Similar changes but in the opposite directions occurred after switching to the dark cycle, resulting to clear rhythmic oscillations during the 24-hour day (Fig 3B-C). Thus, PV’s synaptic excitation and inhibition are modulated in a time-dependent manner during the light/dark cycle.

**Figure 3.**
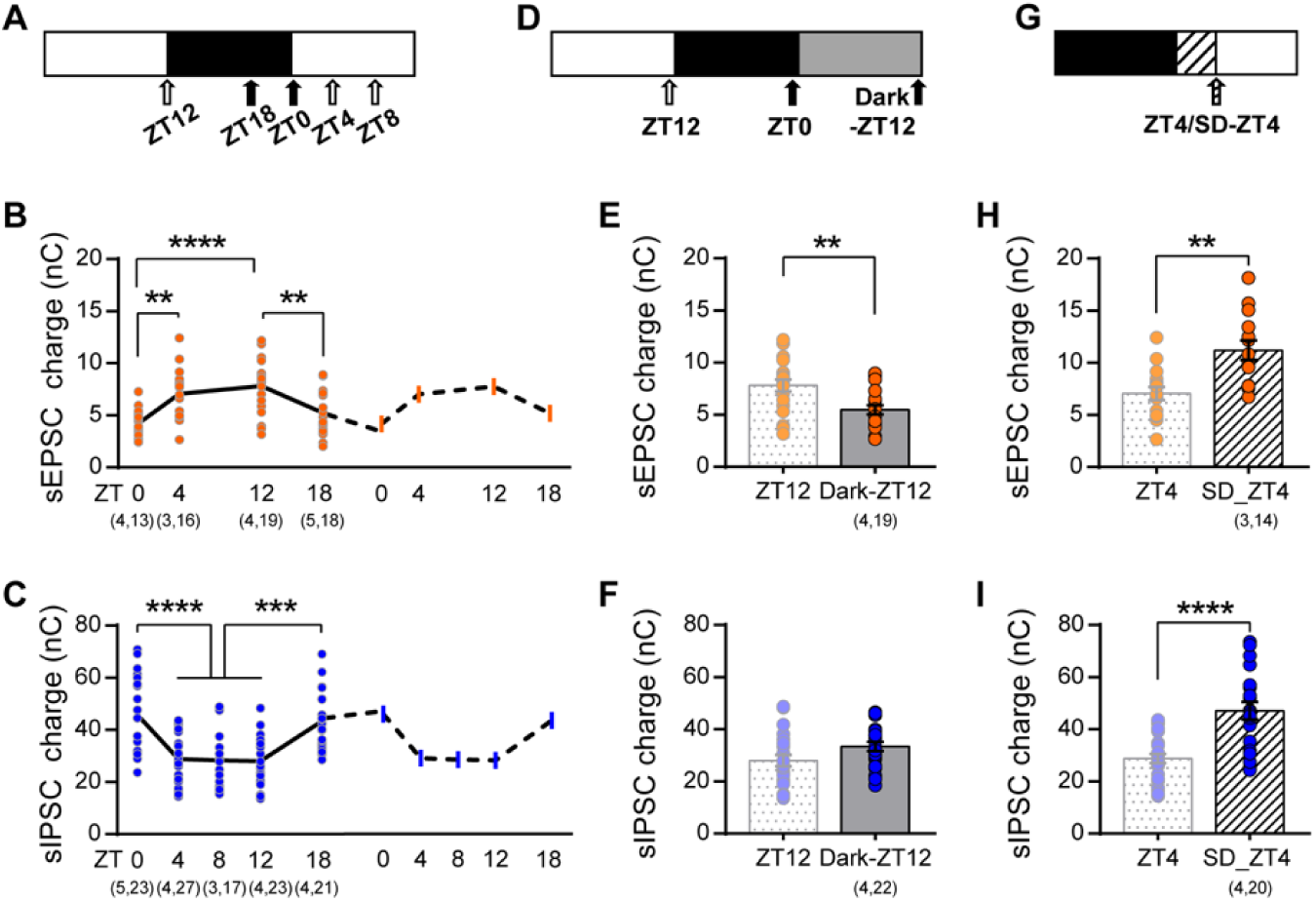
The excitatory and inhibitory synaptic transmission of PV neurons are modulated by experience and sleep. A. Paradigm of experiment. Acute brain slices were obtained at five different times of day (arrowheads). B. Charges of sEPSCs recorded at light phase were higher compared those at dark phase (ZT0: 4.21 ± 0.28 nC; ZT4: 7.05 ± 0.61 nC; ZT12: 7.81 ± 0.59 nC; ZT18: 5.25 ± 0.53 nC; One-way ANOVA F(3,68) = 10.47, P <0.0001, Holm-Sidak post-hoc analysis). C. Charges of sIPSCs oscillate during the light/dark cycle (ZT0: 45.73 ± 2.99 nC; ZT4: 28.85 ± 1.76 nC; ZT8: 28.25 ± 2.77 nC; ZT12: 28.04 ± 2.15 nC; ZT18: 43.37 ± 2.42 nC; One-way ANOVA F(4,106) = 13.65, P <0.0001, Holm-Sidak post-hoc analysis). D. Paradigm of dark exposure. E. Dark exposure prevented the increase in sEPSCs that normally occurs at ZT12 as reported in B (Dark-ZT12: 5.46 ± 0.43 nC; P = 0.0082). F. Dark exposure did not significantly affect the reduction of sIPSCs (ZT12 as reported in C; Dark-ZT12: 33.43 ± 1.74 nC; P = 0.07). G. Paradigm of acute sleep deprivation. H. Mice underwent sleep deprivation further increased the sEPSCs compared to *ad lib sleep* mice (ZT4 as reported in B; SD-ZT4: 11.22 ± 0.92 nC; P = 0.0013). I. Sleep deprivation abolished the reduction of sIPSC (ZT4 as reported in C; SD-ZT4: 47.13 ± 3.39 nC; P < 0.0001).

### Experience- and sleep-dependent regulation of synaptic transmission of PV neurons

We next sought to understand what may drive the opposite changes in excitation and inhibition. Neurons including GABAergic inhibitory neurons in V1 L2/3 are strongly recruited and tuned by direct visual experience(Kerlin, Andermann et al., 2010, Liu, Li et al., 2009). Therefore, visual experience during the light cycle might be involved. To directly test the role of experience, we visually deprived a group of PV:Ai9 mice by dark exposure during the last light cycle prior to experimentation (Dark-ZT12, Fig 3D). Transient visual deprivation during the light cycle abolished the upregulation of sEPSCs (Fig 3E), suggesting that experience plays an important role in regulating excitatory input onto PV neurons. This sort of regulation was not limited to the primary visual cortex. In primary motor cortex (M1), PV neurons showed higher excitatory synaptic transmission by the end of the dark cycle (Fig S2B), during which mice are more active, further supporting the experience-dependency. In striking contrast, we found that the downregulation of IPSC during the light cycle was independent of visual input (Fig 3F), indicative of distinct regulatory mechanisms for synaptic excitation and inhibition during the light/dark cycle. Sleep has been shown to regulate both excitation and inhibition of pyramidal neurons(Bridi et al., 2020, Liu, Faraguna et al., 2010). Therefore, we next studied how acute sleep deprivation (SD) at the beginning of the light cycle may affect the transition of PV’s synaptic transmissions (Fig 3G), during which mice spend around 70% of time sleeping (Fig S2C). SD by gentle handling did not cause obvious increase in corticosterone level (Fig S2E) and its efficacy was confirmed by electroencephalogram (EEG) and electromyogram (EMG) recordings (Fig S2F). SD abolished the reduction in sIPSCs (Fig 3I), indicating that natural sleep is required for the downregulation of synaptic inhibition during the light cycle. Interestingly, SD failed to prevent the increase, and actually further enhanced sEPSCs (Fig 3H), likely owing to an indirect effect by boo sting visual experience. Thus, synaptic excitation and inhibition of PV neurons is mainly regulated by experience and sleep, respectively.

### Acetylcholine regulates inhibitory synaptic properties of PV neurons via M1R

Having established that, in contrast to excitation, PV’s inhibition is modulated in a time- and sleep-dependent manner, we sought to define the mechanism underlying this process. Changes in mIPSC frequency are usually attributed to either altered presynaptic function or synapse number(Han & Stevens, 2009). By examining GAD65-positive presynaptic puncta around the soma of PV neurons (Fig S3A), we found no change in perisomatic inhibitory synaptic density between ZT0 and ZT12 (Fig S3B_1_), but we did see an increase in puncta size at ZT12 (Fig S3B_2_). Positive correlation between presynaptic size and presynaptic function has been reported(Murthy, Schikorski et al., 2001). Therefore, our structural results imply that the inhibitory presynaptic function of PV neurons may be rhythmically modulated.

Several neuromodulators including acetylcholine (ACh) and norepinephrine (NE) have been shown to directly regulate presynaptic properties of different cell types in various brain regions(Choy, Agahari et al., 2018, Kruglikov & Rudy, 2008). Cortical levels of these neuromodulators oscillate during the 24-hour day that correlate with the sleep/wake cycle – high during wake cycle and low during sleep cycle (Jiménez-Capdeville & Dykes, 1993, Kametani & Kawamura, 1991), and they are proposed to be the key mediators of circadian(Frank & Cantera, 2014) and sleep(Tononi & Cirelli, 2014a) in regulating synaptic plasticity. To test if ACh and NE are required for the daily regulation of PV’s inhibitory synaptic transmission (Fig 4A), we first tested the effect of cholinergic and noradrenergic agonists on mIPSCs at ZT12, when mIPSC frequency was low (Fig 2B). Whereas treatment with NE was effectless (Fig S3C), incubation with carbachol (CARB), a non-selective acetylcholine receptor agonist, effectively increased mIPSC frequency (Fig 4B) without changing the amplitude (Fig 4C). This effect was limited to cells recorded at ZT12 but not at ZT0 (Fig 4D-E), indicating that the upregulation of inhibition at ZT0 may utilize overlapping mechanism as cholinergic receptor activation. Interestingly, the effect of ACh seems to be cell type-specific since mIPSCs of pyramidal neurons were insensitive to CARB treatment (Fig S3H). These results suggest that ACh plays a unique role in the daily regulation of PV’s inhibition. To investigate the downstream effector of ACh, we acutely infused brain slices with either muscarinic or nicotinic ACh receptor antagonists after establishing a baseline mIPSC recording at ZT0. Infusion with 50 µM non-selective mAChRs antagonist atropine (Atr) but not nAChRs antagonist (Fig S3F) was sufficient to reduce mIPSC frequency (Fig 4F), which could be rapidly reversed by washing out the drug (Fig S3E_1_). Together, these results suggest the increase in PV’s inhibition by the end of the dark cycle requires mAChRs activation. We further defined the subtype of mAChRs by testing antagonists selective for M1R (pirenzepine, Pir) and M3R (1,1-dimethyl-4-diphenylacetoxypiperidinium iodide, 4-DAMP). Pir showed a quick and reversible effect on mIPSC frequency similar to Atr (Fig 4H) but 4-DAMP had no effect (Fig S3G), suggesting a selective role for M1R. We confirmed these findings by demonstrating that co-incubation with Pir blocked the CARB-induced elevation in mIPSC frequency at ZT12 (Fig 4J). None of the drug treatments affected mIPSC amplitude (Fig 4G, I, K; Fig S3C_2_-G_2_), further supporting the presynaptic locus for this form of modulation.

**Figure 4.**
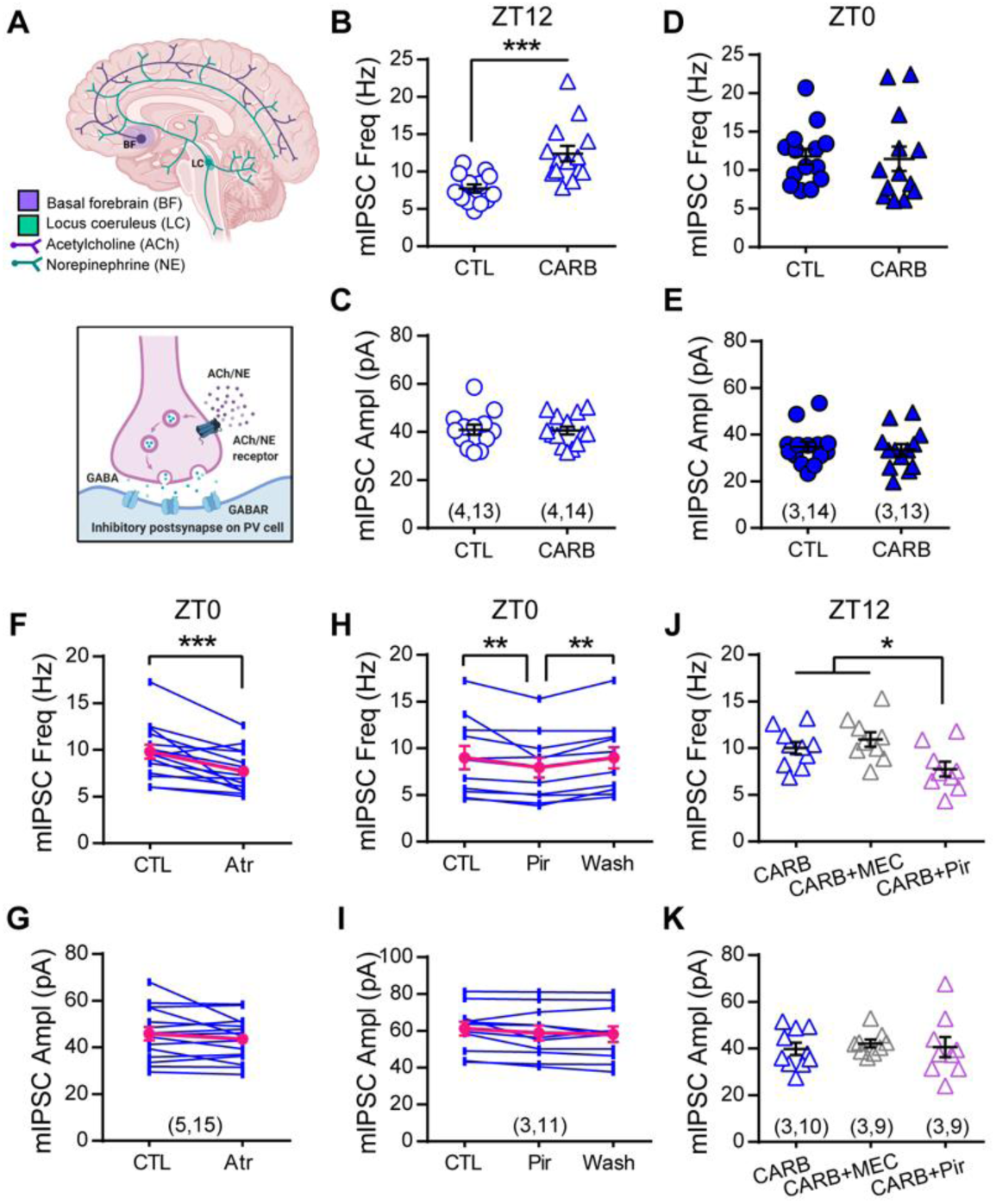
Acetylcholine modulates inhibitory synaptic transmission of PV neurons via M1R during the day. A. Diagram illustrating the potential neuromodulation of synaptic inhibition of PV neurons. B-E. Slices were obtained at ZT0 (B-C) or ZT12 (D-E) and pre-incubated with 50 µM CARB for more than 1h before recording mIPSCs in the presence of drug. Slices from the same animals were used as controls. CARB increased mIPSC frequency at ZT12 (CTL: 7.74 ± 0.55 Hz; CARB: 12.41 ± 1.03 Hz; P = 0.0001), but not ZT0 (CTL:11.79 ± 1.00 Hz; CARB: 11.47 ± 1.60 Hz; P = 0.39). mIPSC amplitude was unchanged at either time (C, ZT12: CTL: 40.94 ± 2.08 pA; CARB: 40.64 ± 1.66 pA; P = 0.99; E, ZT0: CTL: 34.73 ± 2.14 pA; CARB: 33.61 ± 2.39 pA; P = 0.85). F-G. mIPSC frequency (F, CTL: 9.82± 0.74 Hz; Atr: 7.70 ± 0.57 Hz) and amplitude (G, CTL: 45.88 ± 2.87 pA; Atr: 43.64 ± 2.40 pA; P = 0.70) recorded at ZT0 with acute atropine (Atr) wash on. H-I. Effect of acute wash-on and wash-off of pirenzepine (Pir) on mIPSC frequency (H, CTL: 9.00 ± 1.24 Hz; Pir: 7.94 ± 1.07 Hz; Washout: 8.99 ± 1.14 Hz; One-way ANOVA F(2, 20) = 7.483, P = 0.0037) and amplitude (I, CTL: 61.23 ± 3.72 pA; Pir: 58.85 ± 4.06 pA; Washout: 58.25 ± 4.20 pA; One-way ANOVA F(2, 20) = 3.197, P = 0.0624). J-K, Effect of pirenzepine or mecamylamine (MEC) on carbachol-induced elevation in mIPSC frequency at ZT12 (J, CARB: 10.02 ± 0.64 Hz; CARB+MEC: 10.95 ± 0.78 Hz; CARB+Pir: 7.75 ± 0.79 Hz; One-way ANOVA F(2, 25) = 4.848, P = 0.0166) and amplitude (K, CARB: 39.85 ± 2.60 pA; CARB+MEC: 42.17 ± 1.65 pA; CARB+Pir: 40.67 ± 4.34 pA; One-way ANOVA F(2, 25) = 0.1501, P = 0.8614). Sample size is indicated as (mice, cells).

How does activation of M1Rs by ACh alter presynaptic properties? KCNQ potassium channel is a downstream target of M1R activation that influences the membrane potential(Wang & Li, 2016) (Fig S4A) and regulates synaptic release of multiple neurotransmitters including GABA(Martire, Castaldo et al., 2004). By testing the effects of KCNQ2/3 channel opener retigabine (PAM) (Martire et al., 2004) (Fig S4B-C) and blocker XE991 (Fig S4D-E) on PV’s mIPSCs, we found KCNQ is indeed involved in the bidirectional regulation of PV’s inhibition. Furthermore, co-application of XE991 with M1R antagonist Pir completely abolished the Pir-induced reduction in mIPSC frequency recorded at ZT0 (Fig S4F), indicating that KCNQ channels are critical downstream mediators of M1R activation in regulating PV’s inhibition. In summary, we have found that ACh is involved in the daily regulation of PV’s inhibitory synaptic transmission via targeting M1Rs, which then recruit KCNQ channels. This signaling pathway may serve as one important mechanism underlying the daily modulation of PV neuronal function.

### Spontaneous activity of PV neurons oscillates naturally during the light/dark cycle

All our *ex vivo* evidences imply that PV neurons *in vivo* may altered their neural activity at different times of the day. To test this hypothesis, we first performed *in vivo* two-photon calcium imaging to compare PV’s spontaneous Ca^2+^ activity at ZT0 and ZT12 (Fig 5A, Methods). Same group of PV neurons were identified (Fig S5A) and we verified intermittent imaging would not disturb mice’s regular activity cycle (Fig S5H). To avoid visual evoked response and eliminate the effect of arousal state on neural activity, all imaging was conducted in dark with sufficient habituation while mice remained quietly awake and free-running in an air-suspended arena (Fig 5A_3_, Fig S5C). We found that majority of PV neurons had higher spontaneous activity and more events at ZT12 compared to ZT0 (Fig 5B). The mean activity of all PV neurons (indicated by sum(ΔF/F_0_), see method) is significantly larger with right-tilted cumulative distribution at ZT12 (Fig 5C). In addition, neighboring putative pyramidal neurons estimated by their Ca^2+^ response skewness (Dipoppa, Ranson et al., 2018) (Fig S5B) showed opposite change of activity as PVs (Fig 5D-E), which is consistent with higher activity and greater inhibitory output of PV neurons at ZT12 compared to ZT0. These distinct activity patterns are unlikely due to the relatively higher locomotor activity at ZT0 (Fig S5D) since no cross-correlation between neural activity and locomotion was found at either time point (Fig S5E). Therefore, our findings support that PV neurons do alter their spontaneous activity at different times of the day, which influences activity of the surrounding pyramidal neurons.

**Figure 5.**
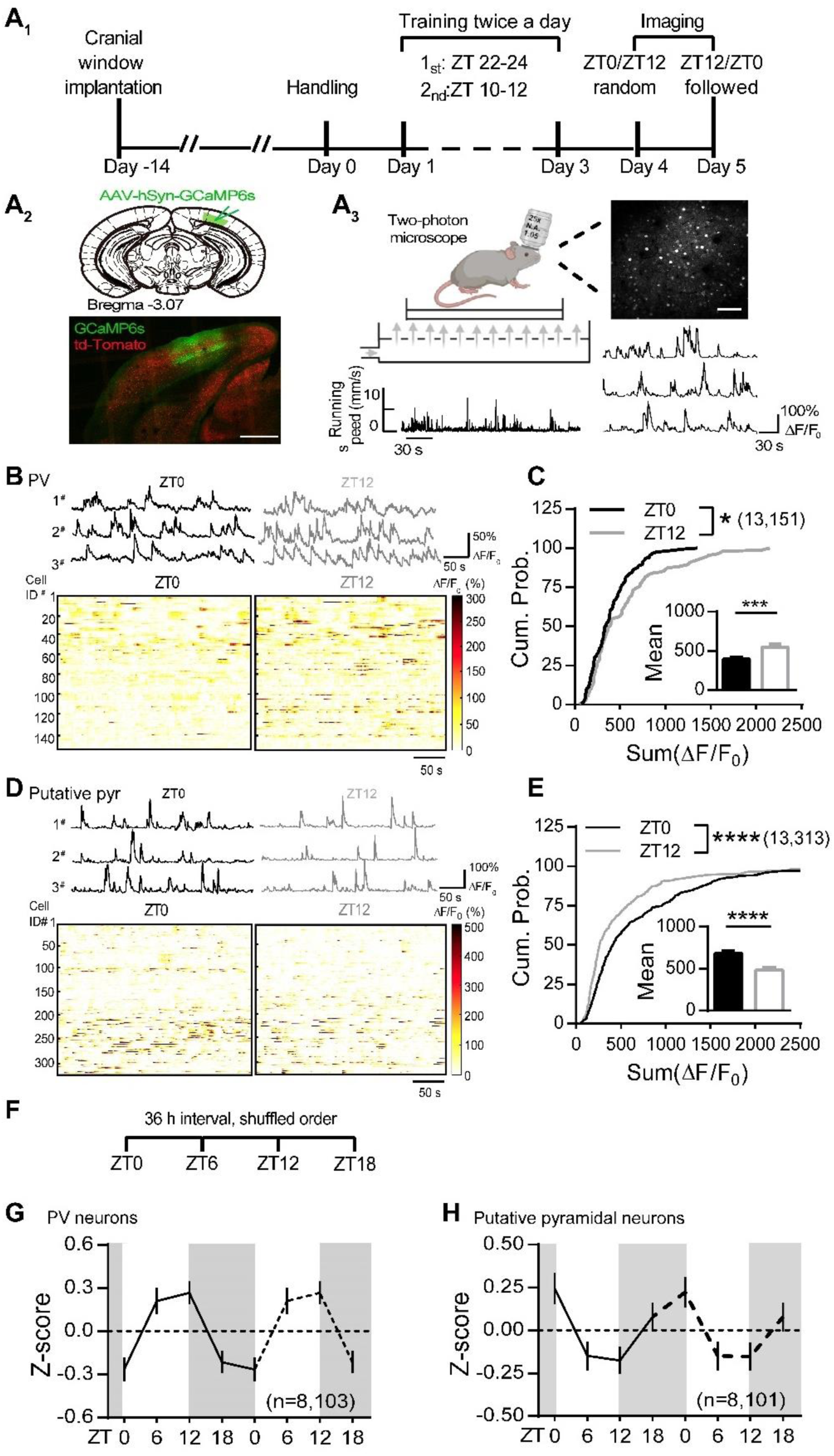
Spontaneous activity of PV neurons oscillates naturally during the light/dark cycle. A_1_. Diagram illustrating the *in vivo* two-photon imaging timeline. A_2_. GCaMP6s (green) was expressed in V1 of the PV:Ai9 mice (red: td-Tomato). Scale: 500 µm. A_3_. Schematics of the calcium imaging showing a mouse that was head-fixed and free running on an air-suspended arena during the imaging section. Locomotor activity was simultaneously monitored (bottom left) along with Ca^2+^ imaging (Top right. Scale: 100 µm). Example spontaneous ΔF/F_0_ traces (bottom right). B. Top panel, Example spontaneous ΔF/F_0_ traces from three PV neurons repetitively imaged at ZT0 and ZT12. Bottom panels, Heatmaps showing the spontaneous ΔF/F_0_ time-series traces of all PV neurons imaged at ZT0 and ZT12. The color code represents the magnitude of ΔF/F_0_(%). C. Cumulative probability distribution of the sum ΔF/F_0_ of all PV neurons imaged at ZT0 and ZT12 (*K-S* test). Insert: the mean sum ΔF/F_0_ (ZT0: 401.6 ± 19.55; ZT12: 559.1 ± 35.29; n =151 cells from 13 mice). D. Top panel, Example spontaneous ΔF/F_0_ traces from three putative pyramidal neurons repetitively imaged at ZT0 and ZT12. Bottom panels, Heatmaps showing the spontaneous ΔF/F_0_ time-series traces of all putative pyramidal neurons imaged at ZT0 and ZT12. E. The cumulative distribution of the sum ΔF/F_0_ of putative pyramidal neurons judged by skewness (*K-S* test, n = 313 cells from 13 mice). Insert, the means of sum ΔF/F_0_. F. Four timepoints repetitive imaging diagram. G-H. Mean activity of PV neurons (G) and putative pyramidal neurons (H) indicated by Z score imaged at four timepoints. The activity patterns were replicated (dotted lines) for another 24-hour to demonstrate the oscillation. *P < 0.05; **P < 0.01; ***P < 0.001; ****P < 0.0001.

The bidirectional oscillations of PV’s synaptic transmission predict changes in PV’s activity *in vivo* are not limited to ZT0 and ZT12. We hence repetitively imaged mice four times in shuffled order at ZT0, 6, 12 and 18 with 36-hour interval (Fig 5F) without interrupting mice’s normal activity cycle (Fig S5H). Consistent with the temporal profiles of sEPSCs and sIPSCs, most PV neurons had higher activity at ZT6 and/or 12 while reduced activity at ZT0 and/or 18 (Fig S5I_1_), leading to an apparent oscillation of the mean activity across the 24-hour day (Fig 5G). In contrast, putative pyramidal neurons mirrored the oscillation (Fig 5H & Fig S5I_2_). Although we only imaged at static timepoints, a daily oscillation of local PV neuronal activity could be deduced from the current result, suggesting a rhythmic regulation of PV neuronal function during the natural light/dark cycle.

### Daily modulation of PV neuronal function correlates with altered dLGN-evoked responses in V1

PV neurons are key players regulating visual response gain. The high and low spontaneous activity of PV neurons that we observed during light and dark cycles, respectively, may be sufficient to alter the response gain. To directly test this, we compared V1 responses evoked by stimulating dorsal lateral geniculate nucleus (dLGN) at ZT0 and ZT12 from the same animals (Fig 6A_1_). We expressed ChR2-mCherry in dLGN neurons and GCaMP6s in V1 L2/3 neurons. An optic fiber was implanted laterally to deliver light stimulation over the dLGN neurons (Fig 6A_2_). 2-3 weeks after surgery, mice were habituated and repetitively imaged in complete darkness using 2-photon microscopy under head-fixation at ZT0 and ZT12. We verified that our light stimulation could not induced retina-evoked response (Fig S6A-B). Then we stimulated dLGN 15 times at 0.1 Hz and recorded evoked Ca^2+^ responses from the same population of neurons at both time points (Fig 6A_3_, see Method). Not all neurons within the FOVs showed dLGN-evoked responses, but those neurons showing responses at either or both time points were counted as the responsive cells and their percentages were calculated for each mouse. As expected, mice had a higher percentage of responsive cells when imaged at ZT0 compared to ZT12 (Fig 6F). The response fidelity, calculated as the proportion of evoked responses out of the 15 dLGN stimulations, in responsive cells showed significantly higher fidelity at ZT0 than ZT12 (Fig 6G). Finally, we compared the amplitude of the dLGN-evoked responses at both time points for each responsive cell and found a bigger response at ZT0 than at ZT12 (Fig 6B-E & 6H), which were inversely correlated with the spontaneous activity of local PV neurons (Fig S6C-D). Together, these results confirm that superficial neurons in V1 have greater evoked responses at ZT0 than at ZT12, which is consistent with weaker PV activity during the dark and stronger activity during the light cycle (Fig S6C-F). Thus, our data suggest that the daily modulation of PV neuronal function is sufficient to directly impact the response gain in the visual cortex.

**Figure 6.**
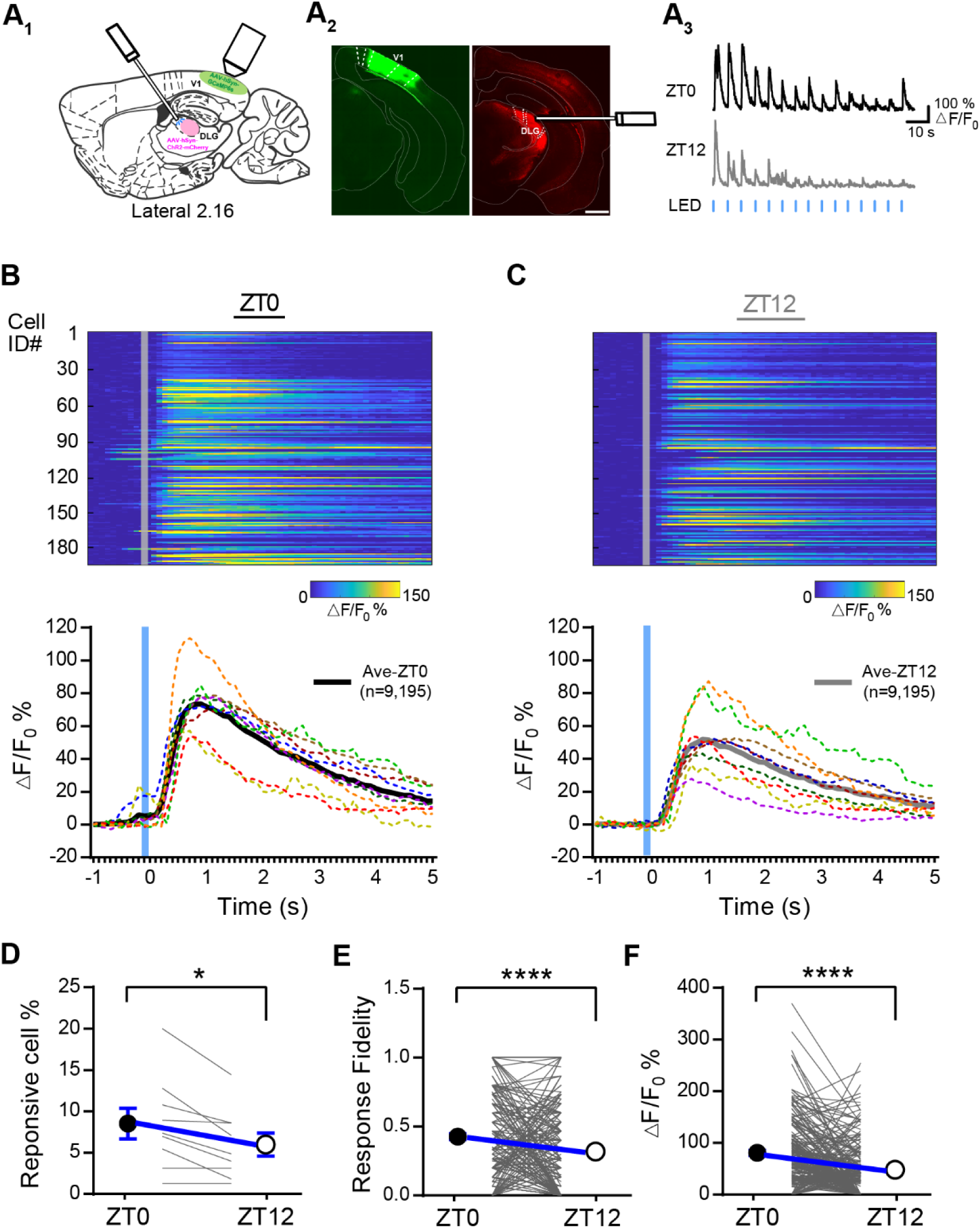
Daily modulation of PV neuronal function correlates with the altered dLGN-evoked responses in V1. A_1_. Schematic showing in vivo dLGN-evoked Ca^2+^ reponses in V1. A_2_. Immunofluorescent staining showing the GCaMP6s (green) and ChR2-mCherry (red) expression in V1 and dLGN respectively. The optical fiber track was also labeled. (Scale: 500 um). A_3_. Response profile of an example neurons for dLGN stimulation during the light/dark cycle. Horizontal blue brief lines represent stimulus frequency. B-C. Top: color-coded calcium signals of all responsive neurons (% ΔF/F_0_). Bottom: The mean Ca^2+^ response waveform for individual mouse (dotted color lines) and all mice (thick black and gray lines). Trials were aligned to LED onset time (gray solid lines). D. The fraction of neurons responsive to dLGN stimulus in ZT0 and ZT12. Each gray line represents one imaging mouse. Black and white circles and error bars represent the mean and SEM (ZT0: 8.51 ± 1.86; ZT12: 5.96 ± 1.39; n = 9 mice). E. The Response fidelity of each cell in ZT0 and ZT12. Black and white circles and error bars represent the mean and SEM (ZT0: 0.42 ± 0.022; ZT12: 0.31 ± 0.023; n = 195 cell). F. Comparison of peak evoked response of each cell in ZT0 and ZT12. Black and white circles and error bars represent the mean and SEM (ZT0: 81.01 ± 4.79; ZT12: 48.42 ± 3.82; n = 195 cell).

## Discussion

Proper function of PV neurons is critical for maintaining cognitive performance and highly relevant with a variety of brain disorders. Here we report that PV neurons in V1 are subjected to tight modulation during the natural light/dark cycle in a time- and sleep-dependent manner. Acetylcholine and its downstream M1R are critically involved in mediating this regulation. Changes in PV’s activity *in vivo* negatively correlate with the activity pattern of surrounding pyramidal neurons and dLGN-evoked responses in V1, supporting the physiological significance of such daily modulation. Therefore, our study unveils a novel regulatory mechanism for PV neurons, provides implications on how daily rhythm and sleep might modulate the brain function.

The inhibitory connections between PV and pyramidal neurons in cortices are known to be developmentally regulated in an experience-dependent manner(Jiang, Huang et al., 2005). However, in mature brain, inhibitory synapses become more stable and are resistant to experience deprivation(Gao, Whitt et al., 2017). The lack of cortical iLTD in adult brain(Huang, Gu et al., 2010) suggests that these synapses are strongly anchored. Surprisingly, we found that the efficiency of the inhibitory connection between PV and pyramidal neurons dramatically differs between the light and dark cycle, consistent with our previous finding showing the up- and down-regulation of mIPSC frequency of cortical pyramidal neurons(Bridi et al., 2020). These results imply that inhibitory synapses in adult brain are not as rigid as previously thought, but instead are flexibly modulated during the 24-hour day. Since inhibition is critical for cortical plasticity, it will be intriguing to investigate whether cortical plasticity may be differentially expressed during day.

The bidirectional changes in PV’s excitatory and inhibitory synaptic transmission show strong time dependency. Both of them rapidly transit from high to low or vice versa after entering the next cycle, and then they remain relatively stable throughout the rest of the cycle regardless of similar sleep/wake activity (Fig S2C), suggesting a ‘Clock’-dependent regulation. Circadian has long been proposed to regulate synaptic efficacy, especially the excitatory synapses of excitatory neurons (Frank & Cantera, 2014). We previously found that the inhibitory synaptic strength of cortical pyramidal neurons also shows rhythmicity(Bridi et al., 2020). Hence, inhibitory synaptic efficacy of both excitatory and inhibitory neurons is likely under circadian modulation as excitatory synapses. In addition to circadian, many evidences support the role of sleep in regulating excitatory synaptic plasticity(Tononi & Cirelli, 2020). Indeed, we found that sleep is essential for regulating the transition of PV’s inhibitory synaptic transmission during the early light cycle. Therefore, the daily modulation of PV’s synaptic transmission is dependent on both ‘Clock’ and sleep. We further identified that ACh via M1R activation is critical signal mediating the bidirectional modulation of PV’s inhibitory synaptic efficacy. Interestingly, the working dynamic of the same cholinergic signaling seems to be different for up- and downregulation of PV’s mIPSCs. High mIPSCs at ZT0 can be rapidly reduced by blocking the ACh-M1R signal (Fig 4F&H), while low mIPSCs at ZT12 requires prolong activation of this pathway (Fig 4B & Fig S3D). This distinction suggests that chronic high level of cortical ACh is critical for maintaining the inhibitory synaptic efficacy during the dark cycle. However, these inhibitory synapses are more resistant to abrupt increase in ACh, such as that reported in wake and rapid eye movement (REM) sleep(Ma, Zhang et al., 2020, Marrosu, Portas et al., 1995), when cholinergic tone is low during the late light cycle. Neuromodulators are candidates mediating both the sleep- and clock-dependent synaptic regulation(Frank & Cantera, 2014, Tononi & Cirelli, 2014b). The circadian rhythm of cortical ACh level(Jiménez-Capdeville & Dykes, 1993, Kametani & Kawamura, 1991) well correlates with the daily modulation of PV’s inhibitory synapses, which can account for both its sleep- and clock-dependency. Thus, our results provide evidences support the active role of ACh in the daily regulation of the inhibitory synapses specifically of PV neurons.

It is surprising to find that the ACh-dependent regulation of mIPSCs is unique for PV but not pyramidal neurons in V1 L2/3 (Fig S3H) even though they receive inhibition from overlapping resources(Harris & Mrsic-Flogel, 2013). This might be due to the unique enrichment of M1Rs at presynaptic terminals targeting PV neurons. The diversity of subtypes and subcellular distribution of neuromodulator receptors have been long known(Dembrow & Johnston, 2014, Palomero-Gallagher & Zilles, 2019). For example, it has been shown that nAChR activation selectively potentiates the thalamocortical but not the intracortical synapses of L3 pyramidal neurons in the primary somatosensory cortex(Gil, Connors et al., 1997). A recent study reported that the NDNF interneurons in V1 L1 specifically target supragranular PV neurons(Cohen-Kashi Malina, Tsivourakis et al., 2021), which could be another explanation for the selectivity of ACh. On the other hand, we have not examined other neuromodulators such as serotonin and histamine, both of which have dense innervation in the visual cortex and also show rhythmic brain levels(Kosofsky, Molliver et al., 1984, Manning, Wilson et al., 1996). Their potential contribution cannot be excluded although our results do suggest manipulating cholinergic signaling alone is sufficient to bidirectionally alter PV’s mIPSCs.

PV’s opposite oscillations in E and I temporally coincide with their spontaneous activity in vivo, which is congruent with the role of E/I ratio in controlling somatic firing(Gidon & Segev, 2012). Therefore, PV’s spontaneous activity likely shares similar time- and sleep-dependent regulation during the day and the circadian rhythm of the cholinergic signaling may also be involved. This provides one explanation to reconcile the discrepancy regarding the role of acute sleep stages and/or sleep history in modulating neuronal firing(Grosmark, Mizuseki et al., 2012, Hengen et al., 2013, Miyawaki & Diba, 2016, Mizuseki & Buzsaki, 2013, Niethard et al., 2016, Vyazovskiy et al., 2009, Watson, Levenstein et al., 2016). When studies are conducted during the early light cycle, sleep and sleep history indeed alter neuronal firing(Grosmark et al., 2012, Miyawaki & Diba, 2016, Niethard et al., 2016, Vyazovskiy et al., 2009, Watson et al., 2016), probably due the sleep-dependent changes in cortical levels of ACh as well as other neuromodulators(Bridi et al., 2020, Hasselmo, 1999, Nadim & Bucher, 2014), and neuronal properties such as synaptic efficacy have not yet reached steady state. For others examining neuronal firing during the late light cycle when the tone of most neuromodulators remains low and the neuronal properties are already stable, switching between sleep stages or even sleep history may not be sufficient to cause changes(Hengen et al., 2013, Mizuseki & Buzsaki, 2013).

Our results suggest different mean activity between the light and dark cycle for PV and putative pyramidal neurons, which seems to be contradictory to previous finding showing similar mean firing rate of regular spiking neurons between the cycles(Hengen et al., 2016). This might be due to sampling bias. While we only studied neurons in supragranular layer at static timepoints, their multiunit recording usually targeted neurons in deeper cortical layers over long period of time. It is necessary to point out that the mean activity of non-PV non-putative pyramidal neurons are not different between ZT0 and ZT12 (Fig S5F&G), further suggesting cell type-specific dynamic of neural activity. In addition, we have only shown time-dependent changes in neural activity when mice were quietly awake, which may partially reflect the effect of sleep history but not acute sleep stages. PV neurons in somatosensory cortex are reported to fire at higher rate at wake and REM sleep compared to NREM sleep during the early light cycle(Niethard et al., 2016). How PV neurons in V1 are modulated by acute sleep stages and how PV neurons in other cortical regions are modulated daily remains to be examined.

PV neurons play a critical role in gain control. Their daily oscillation in neural activity may cause the neural network where they reside in to respond differentially to their inputs(Ferguson & Cardin, 2020), which may underlie the daily fluctuation in cognitive performance(Muto, Jaspar et al., 2016). Studies in the visual cortex have provided solid evidences to show that PV activity determines the gain of visual response(Wilson et al., 2012, Zhu et al., 2015). Although the magnitude of PV oscillation is not as great as the optogenetic manipulation used previously(Atallah et al., 2012), they may be sufficient to alter the response gain in V1. Indeed, the dLGN-evoked responses in V1 are significantly different between ZT0 and ZT12 following the gain rule of PV neurons, strongly suggesting that daily modulation of PV neuronal function is physiologically important. In the visual circuit, light sensitivity of retina has long been known to show circadian rhythm(Bassi & Powers, 1986) and arousal state influences the retina output(Schroder, Steinmetz et al., 2020). Here we directly activated dLGN neurons to mimic visual input, therefore, we could exclude the potential contribution of retina to the alternating V1 responses.

Additionally, our findings also provide some mechanistic implications for neuronal dysfunction in brain diseases such as Alzheimer’s disease (AD). Impaired PV neuronal function is considered a key abnormality in AD brains(Cattaud, Bezzina et al., 2018, Verret et al., 2012), which is linked to reduced gamma oscillation and impaired cognitive performance(Palop & Mucke, 2016, Verret et al., 2012). Meanwhile, disturbed sleep and circadian rhythm and impaired cholinergic function are commonly observed in AD patients, which could be initiated decades before the diagnosis and are recognized as risk factors for AD(Falgas, Walsh et al., 2021). Here by showing how synaptic and neuronal properties of PV neurons are rhythmically regulated, we provide a potential explanation and open new research directions to study disease-associated PV impairment.

## Materials and Methods

### Animals

All experimental procedures were approved by the Institutional Animal Care and Use Committees at the Interdisciplinary Research Center on Biology and Chemistry, Chinese Academy of Science. Homozygous PV-ires-cre mice (B6;129P2-*Pvalb*^*tm1(cre)Arbr*^/J, PV:cre, Jax Laboratory) were used in this study. They were either crossed with Ai9 mice (B6.Cg-*Gt(ROSA)26Sor*^*tm9(CAG-tdTomato)Hze*^/J, Jax Laboratory) to genetically label PV neurons with tdTomato (PV:Ai9), or with Ai32 mice (B6;129S-*Gt(ROSA)26Sor*^*tm32(CAG-COP4*H134R/EYFP)Hze*^/J, Jax Laboratory) to express channelrhodopsin-2 (ChR2) and EYPF specifically in PV neurons (PV:Ai32). 5-10 weeks old male mice were used only. All mice were housed with a 12 h:12 h light/dark cycle with *ad libitum* access to food and water. Mice with shifted light/dark cycle were entrained in customized entrainment chamber for at least 2 weeks before experiments. The light-on time (ZT0) was adjusted according to the experiment.

### Acute Brain Slices Preparation

300 µm thickness acute coronal brain slices containing the primary visual cortex or motor cortex were prepared as described previously (23). Briefly, mice were anesthetized with isoflurane vapor and rapidly sacrificed within 2h before the light on (ZT0) or light off (ZT12). Slices were cut in ice-cold dissection buffer containing the following: 212.7 mM sucrose, 5 mM KCl, 1.25 mM NaH_2_PO4, 10 mM MgCl_2_, 0.5 mM CaCl_2_, 26 mM NaHCO_3_, and 10 mM dextrose, bubbled with 95% O_2_/5% CO_2_. Slices were transferred and incubated at 30 ºC for 30 min with artificial cerebrospinal fluid (ACSF) containing 119 mM NaCl, 5 mM KCl, 1.25 mM NaH_2_PO_4_, 1 mM MgCl_2_, 2 mM CaCl_2_, 26 mM NaHCO_3_, and 10 mM dextrose, saturated with 95% O_2_ and 5% CO_2_. After that, slices were maintained at room temperature (RT) for at least 30 min before recording.

### Whole-Cell Recording

Whole-cell recordings were made from the tdTomato-expressed PV neurons in layer 2/3 of V1 or M1. Slices were submerged in the recording chamber perfused with 30 ± 0.5 ºC perfusion buffer at 2 ml/min. Cells were visualized by an upright fluorescence microscope (Olympus XT640-W). Borosilicate glass pipette recording electrodes with 3-5 MΩ resistance were filled with different internal solutions according to the experiment, all of which were adjusted to pH 7.3-7.4, 285-300 mOsm. Data were filtered at 2kHz and digitized at 10 kHz using Digidata 1550A (Molecular Devices, CA, USA). Only cells with input resistant Ri ≥ 100 MΩ and access resistance Ra ≤ 25 MΩ were used for analysis. In addition, cells were discarded if Ri and Ra changed more than 25% during the recording.

#### a) Miniature postsynaptic current recordings

For mEPSC recordings, 1 μM TTX, 100 μM DL-APV and 20 μM gabazine were added to the ACSF to isolate AMPAR-mediated mEPSCs. The internal solution contained 8 mM KCl, 125 mM cesium gluconate, 10 mM HEPES, 1 mM EGTA, 4 mM MgATP,0.5 mM NaGTP, 10 mM Na-phosphocreatine, and 5 mM QX-314. Cells were held at -80 mV.

For mIPSC recordings, 1 μM TTX, 100 μM DL-APV and 20 μM CNQX were added into the bath. The internal solution contained 8 mM NaCl, 120 mM cesium chloride, 10 mM HEPES,2 mM EGTA,4 mM MgATP,0.5 mM NaGTP, and 10 mM QX-314. Cells were held at -80 mV for experiments in Figure 2 and -60 mV for those in Fig 4 and Fig S4.

#### b) Spontaneous postsynaptic current recordings

Spontaneous EPSCs were recorded in regular ACSF without any blocker. Cells were held at -55 mV, the measured reversal potential of the inhibitory current (Fig S2A). To record spontaneous IPSCs, ACSF containing 100 uM DL-APV and 20 uM CNQX was used and cells were held at -60 mV during the recording. Internal solution for sEPSC and sIPSC was similar to those used for mEPSC and mIPSC except no QX-314 was added.

#### c) ChR2-assisted mapping of the connectivity between PV and pyramidal neurons

Acute brain slices containing V1 from PV:Ai32 mice were prepared. L2/3 pyramidal neurons were identified by their morphologies and voltage clamped at -50 mV with the same internal solution and perfusion buffer as mIPSCs recording. A 0.45 mm^2^ square region surrounding the recorded neuron was divided into 15 by 15 grids. Brief blue light pulses (10 ms at 100% intensity with 1 s inter-stimulus interval) were controlled by Polygon400 (Mightex, USA) and delivered via a water immersion objective (10x, 0.30 NA, Olympus) to each grid in a fixed pseudo-random order and repeated 3 times. Light-evoked inhibitory postsynaptic currents (_LE_IPSC) were recorded.

### Sleep deprivation

Mice were kept awake by gentle handling, during which EEG/EMG signals were acquired simultaneously.

### Visual deprivation

PV:Ai9 mice entrained to the regular light/dark cycle were allowed to habituate to the customized light-proof, well-ventilated entrainment chambers in their homecages at least 2 days before the experiment. Visual deprivation was achieved by keeping one group of mice in the dark during the last light cycle before they were sacrificed for experiment.

### Virus injection and cranial window preparation

6-8 week old PV:Ai9 mice were maintained anesthetized by 1-2% isoflurane vapor and head-fixed in a stereotaxic frame. Ophthalmic ointment (Cisen, China) was applied to the eyes and mice were kept on a heating pad where the temperature was maintained at around 37 ºC. Skin was sterilized by iodine and a craniotomy was made above the right visual cortex. A 3×3 mm circular piece of skull was removed (3.0 mm post the bregma suture, 2.0 mm from the mild line suture) and the dura was kept intact and moist with sterile saline. 50-100 nl AAV2-hSyn-GCaMP6s (1×10^12^ genome copies per ml, Taitool, China) was delivered using a bevelled glass pipette by a microsyringe pump (Stoelting, USA) at 30 nl/min. Total 4-5 injections were made at a depth of ∼ 250 µm in the primary visual cortex. After the virus injection, a cover glass was glued to the skull over the circular opening by using polyacrylic glue (Elmer’s, USA). A customized headplate was then cemented to the skull around the center of the cranial window. Skin was sutured back around the headplate and lidocaine (Sigma, USA) and triple antibiotic ointment (DearGo, China) were applied around the wound. Mice were transferred back to their homecages after fully recovered from anesthesia and were closely monitored during the following week. Virus was allowed to expressed for at least 14 days before the experiment.

### In vivo two-photon calcium imaging

The whole procedure was illustrated in Figure 1A. Specifically, all mice were gentle handled 5-10 min for 2-3 times between ZT0-4 on day 0. From day 1 to 3, mice were allowed to habituate to the head fixation and free running in the mobile homecage (Neurotar, Finland) under the 2-photon microscopy twice a day, 1 hour each between ZT10-12 and ZT22-24. All mice were imaged at least twice (ZT0 and ZT12) post training. Some mice were repeatedly imaged 4 times at ZT0, ZT6, ZT12, and ZT18 with random order. All imaging sections were spaced out with at least 36 hours interval, during which mice were housed in their homecages with activity monitored by video.

For each imaging section, mice were head-fixed under the microcopy at least 10 min before the imaging started. Imaging was performed using a two-photon laser-scanning microscope (FVMPE-RS, OLYMPUS) coupled with a Mai Tai Insight DS Dual-OL laser (Spectra Physics). The tunable laser was set to 920 nm (∼30-70mW average power on the sample) for GCaMP6s and the 1040 nm laser was used for exciting the tdTomato. The emission filter set consisted of a 570 nm dichroic mirror and a 540/50 nm band-pass filter. Fluorescence was detected using GaAsP photomultiplier tubes. Images of 512 × 512 pixels (x/y resolution: 0.663 um/pixel) were scanned in a line mode repeated 3 times by a resonant galvanometer and 3000 to 6000 frames were acquired at 10 Hz for each field of view (FOV). The locomotor activity of all mice during all imaging section was monitored with Motion Tracker (Neurotar, Finland) and/or camera.

In order to image the same population of neurons across sections, the same FOV were tracked by vascular features, cortical depth and spatial location of landmarks (bright and stable structures of somas). Only neurons showing clear and similar cell morphology were included for analysis.

### dLGN optogenetic manipulations

500 nl AAV2-hSyn-hChR2(H134R)-mCherry virus or AAV2-hSyn-mCherry-WPRE (Taitool) was injected into the right dLGN with stereotaxic atlas (AP -2.3; ML 2.15; DV 2.5; <90°) at a rate of 50 nl/min. A 200 µm OD optical fiber (NA = 0.37, Inper) was implanted lateral over dLGN with the same coordination. Cranial window over the right primary visual cortex was prepared and AAV virus expressing GCaMP6s was injected into the superficial layer of V1 as describe above. The recovery and habituation were similar as the regular 2-photon imaging. Optogenetic stimulation of dLGN was achieved using a 470 nm Fiber-Coupled LED (Thorlabs, M470F3) controlled by a LED Driver (Thorlabs, DC2200). Black ceramic sleeve was used to prevent the light form leaking. Light intensity at the fiber tip was set at 75-80 mW/mm^2^ and 10 ms light pulses were delivered at 0.1 Hz for 15 trials. The GCaMP6s signal was acquired in the same way as described in the above section. At the end of experiment, animals were sacrificed and 50 µm coronal brain slices were sectioned with a vibratome to confirm the position of optic fiber an ChR2 expression (Fig 6A_2_).

### Quantification and Statistical Analysis

#### Whole-cell recordings

##### a) Miniature postsynaptic currents

mEPSCs and mIPSCs were analyzed using the MiniAnalysis program (Synaptosoft, Decatur, GA) as described previously (Michelle C.D. et al.,2020). Event detection threshold was set at 3 times over the RMS noise. At least 300 events with rise time ≤ 3 msec for mEPSCs and ≤ 5 msec for mIPSCs were selected for each cell to calculate frequency and amplitude. And non-overlapping events were used to construct the average trace and estimate the decay time.

##### b) Spontaneous postsynaptic currents

Spontaneous EPSCs and IPSCs were analyzed by calculating the unit charge (nA/s) with customized code (MATLAB, MathWorks) as described previously (Michelle C.D. et al.,2020). Briefly, the root mean square (RMS) noise of each 500 msec segment was calculated and subtracted. Charge was calculated as the integral of the post-subtracted signal of total 3-4 min of recording.

##### c) ChR2-assisted mapping of the connectivity between PV and pyramidal neurons

The max amplitude of the response evoked within 30 msec time window post light stimulation was measured. Only reliably evoked responses (2 out of 3 repeats) larger than 60 pA were considered as _LE_IPSCs and the mean was used as the measurement of the connection strength. A heatmap showing the connection strength for each recorded neuron was generated. Maps from each group were either aligned by the patched cell soma (soma-align) or slice pia (pia-align). For soma alignment, maps from the same group were simply averaged since the relative soma position of the recorded neuron was the same across maps. For pia alignment, the 369×680 pixels DIC image showing the shape of the cortical slice, the 15 by 15 grids, and the position of the recording electrode using the 10x objective was taken at the end of each recording. The variation in cortical thickness was first corrected by scaling individual DIC image to match that in the reference image (ref, Figure 1A_3_ top right). The amplitude of the _LE_IPSCs was used to define the pixel values for the corresponding grid in the scaled image. Then the pia in the ref was aligned with that of the scaled DIC image and the pixel value was projected onto the ref. After projecting _LE_IPSCs from all neurons, the average was calculated for each pixel in the reference image and the pia-aligned map was generated and compared between ZT0 and ZT12. All analyses were done by Clampfit (Molecular Devices, CA, USA) and customized matlab programs.

#### Calcium image analysis

Images were analyzed *post-hoc* using customized matlab programs modified based on the original open-source package available online (https://github.com/flatironinstitute/CalmAn-MATLAB). Briefly, stacked time-series images were registered by aligning to a target image using a cross-correlation-based registration algorithm (discrete Fourier transformation, DFT, algorithm) to correct brain motion.(Xin, Zhong et al., 2019) The target image was obtained by mean projection of visually identified frames (≥ 200 frames) with few motion artifacts. Regions of interest (ROIs) were drawn for individual neuron manually based on neuronal shape using a customized GUI and the raw fluorescence signal (F) for all ROIs from all frames was extracted. To calculate ΔF/F_0_, the initial 200 frames for each cell were excluded to avoid imaging artifact.

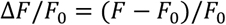

Baseline fluorescence (F_0_) was computed via a sliding-window (400 frames for PV neurons and 550 frames for non-PV neurons) using a 30% quantile cut-off. The correlation coefficient (R^2^) between the converted ΔF/F_0_ waveform and the raw F waveform was calculated and only cells with R^2^ larger than 0.70 were included for final analysis. To estimate neural activity from the calcium signals, the sum of ΔF/F_0_ (sum(ΔF/F_0_)) was calculated after subtracting the 3 times of the minimal root mean square (RMS) noise of each cell.

#### Light-evoked Ca^2+^ signal analysis

The dLGN-evoked Ca^2+^ responses in the superficial layer of V1 was calculated as (F-F_0_)/F_0_ x 100%. The average of the 10 frames (1s) before the onset of light stimulation was used as F_0_. The peak Ca^2+^ intensity within the 1 s time window following stimulus onset was measured as the F. Only responses larger than 15% were included for final analysis. To analyze the fraction of responsive neurons, neurons showing at least 1 effective response at either ZT0 or 12 were counted and divided to the total neurons that had detectable spontaneous activity within the FOV. Response fidelity for each responsive neuron was calculated as the proportion of evoked responses out of the 15 stimuli.

### Statistics

For all figures, the sample size is indicated as (mice, cells). Statistical analysis was performed by Prism V6.0 software (GraphPad Software, Inc.). For two groups comparison, the Wilcoxon rank-sum test and Mann-Whitney test were used for paired and unpaired data respectively. For more than two groups analysis, the ordinary one-way ANOVA followed by Holm-Sidak post hoc test was used. Cumulative distributions were compared with the Kolmogorov-Smirnov (KS) test. Error bars in all figures indicate standard error of mean (s.e.m). Only significant comparisons were labeled in the figures. The level of significance was set at *P* < 0.05. \**P* < 0.05; \*\**P* < 0.01; \*\*\**P* < 0.001; \*\*\*\**P*<0.0001.

## Data and Code Availability

The customized MATLAB codes are available on GitHub.

## Acknowledgments

We thank Dr. Junying Yuan, Dr. Alfredo Kirkwood, Dr. Aaron Gitler for insightful discussion. This work is supported by NSFC (Grant No. 32070963, 31700917), Shanghai Science and Technology Development Funds (Grant No. 22ZR1475100), and Shanghai Municipal Science and Technology Major Project (Grant No. 2019SHZDZX02) to K.-W.H..

## Author Contributions

Conceptualization, K.-W.H.; Methodology, F.-J.Z., X.-T.Z., X.M. and K.-W.H.; Investigation, F.-J.Z., X.-T.Z. & Y. L.; Data Analysis, F.-J.Z., X.-T.Z., Y.Z. and K.-W.H.; Writing, F.-J.Z. and K.-W.H.; Funding Acquisition, K.-W.H; Supervision, K.-W.H..

